# piRNAs regulate a Hedgehog germline-to-soma pro-aging signal

**DOI:** 10.1101/2021.08.09.455665

**Authors:** Cheng Shi, Coleen T. Murphy

## Abstract

The reproductive system regulates the aging of the soma through competing anti- and pro-aging signals. Germline removal extends somatic lifespan through conserved pathways including Insulin, mTOR, and steroid signaling, while germline hyperactivity cuts lifespan short through mechanisms that remain elusive. Here, we show that mating-induced germline hyperactivity leads to the dramatic downregulation of piRNAs, which in turn releases silencing of their targets, including the Hedgehog-like ligand encoding genes *wrt-1* and *wrt-10*, ultimately causing somatic collapse and early death. Germline-produced Hedgehog signals require PTR-6 and PTR-16 receptors for mating-induced body shrinking and lifespan shortening. Our results reveal an unconventional role of the piRNA pathway in transcriptional regulation of Hedgehog signaling, as well as a new role of Hedgehog signaling in the regulation of longevity and somatic maintenance. Our data suggest that Hedgehog signaling is controlled by the tunable piRNA pathway to encode the previously unknown germline-to-soma pro-aging signal. Mating-induced downregulation of piRNAs in the germline and subsequent signaling to the soma via the Hedgehog pathway enables the animal to tune its somatic resource allocation in response to germline needs to optimize reproductive timing and survival.

## Introduction

Longevity is plastic, and is influenced by both external (e.g., diet) and internal factors, such as reproductive demands. Communication between the germline and soma allows animals to prepare a coordinated response to various physiological and environmental challenges. Animals couple nutrient availability to reproduction (Wade and Schneider, 1992), a typical example of soma-to-germline communication, through conserved signaling pathways, including the Insulin, AMPK, and mTOR pathways (Chantranupong et al., 2015, Dupont et al., 2014, Efeyan et al., 2015, Templeman and Murphy, 2018, Laws and Drummond-Barbosa, 2017). Conversely, the status of the germline profoundly influences somatic tissues: germline removal extends lifespan (Hsin and Kenyon, 1999; Arantes-Oliveira et al., 2002; Flatt et al., 2008; Min et al., 2012), while germline hyperactivity decreases lifespan and leads to dramatic changes in somatic physiology in animals across great evolutionary distances (Shi and Murphy, 2014, Shi et al., 2017, Shi and Murphy, 2021).

*Caenorhabditis elegans* has been a useful model to identify mechanisms underlying germline-soma communication. Ablation or removal of the *C. elegans* germline significantly extends lifespan, increases fat accumulation, and enhances resistance to various stresses (Hsin and Kenyon, 1999, Arantes-Oliveira et al., 2002, Ben-Zvi et al., 2009, O’Rourke et al., 2009, Alper et al., 2010, Vilchez et al., 2012). However, concomitant removal of the somatic gonad eliminates lifespan extension and other somatic benefits, suggesting the existence of two opposing signals: a germline pro-aging signal and a somatic gonad pro-longevity signal (Hsin and Kenyon, 1999, Lapierre and Hansen, 2012). The somatic gonad pro-longevity signal pathway has been characterized using germlineless worms (Berman and Kenyon, 2006, McCormick et al., 2012, Goudeau et al., 2011, Lapierre et al., 2011, Antebi, 2013, O’Rourke and Ruvkun, 2013, Ratnappan et al., 2014, Boulias and Horvitz, 2012, Kenyon). Dafachronic acids are thought to be the most upstream pro-longevity signal from the somatic gonad (Yamawaki et al., 2010, Antebi, 2013). Insulin, mTOR, and steroid signaling are required in the soma for germline loss-mediated lifespan regulation, suggesting they mediate the somatic gonad prolongevity effect (Berman and Kenyon, 2006, Gerisch et al., 2007, Flatt et al., 2008, Lapierre and Hansen, 2012, McCormick et al., 2012). By contrast, the identity of the initial pro-aging signal originating from the germline remains elusive. Identifying this pro-aging signal is critical for understanding how animals tune their aging rates in response to germline activity.

While most studies of the germline’s influence on lifespan have compared germlineless to intact, unmated animals (Hsin and Kenyon, 1999, Lapierre and Hansen, 2012), less is known about the influence of the hyperactive germline on longevity. Mating significantly accelerates germline proliferation and ultimately leads to somatic collapse and early death (Shi and Murphy, 2014, Shi et al., 2017), suggesting that the germline pro-aging signal is significantly amplified by mating. Therefore, mated worms are an ideal system in which to identify the mysterious germline pro-aging signal.

Here we set out to identify the underlying mechanism of hyperactive germline-induced shrinking and early death. Our transcriptional analyses of isolated germlines revealed that a specific subset of piRNAs is downregulated in response to mating, de-repressing Hedgehog-related genes. Hedgehog signaling communicates the status of the germline to somatic cells, resulting in mating-induced shrinking and early death. Thus, somatic responses to germline hyperactivity are tuned by piRNA regulation of a major developmental signaling pathway.

## Results

### Mating induces significant transcriptional changes in the germline

Mating leads to body shrinking and decreased lifespan (Shi and Murphy, 2014). Day 1 adult spermless hermaphrodites (*fog-2*) that are mated for 24hrs with Day 1 adult males live 40% shorter than their unmated counterparts, and they also shrink by up to 30% (**Figure 1A-B, Figure S1A**). The longevity decrease after mating is mediated by multiple factors including seminal fluid transfer, male sperm-induced germline hyperactivity, and male pheromone toxicity (Shi and Murphy, 2014, Maures et al., 2014, Shi et al., 2017, Shi et al., 2019), among which only the mechanism underlying seminal fluid-mediated death has been characterized; this involves insulin-like peptides and the insulin/FOXO signaling pathway. The 30% shrinking of the adult body size is a germline-specific pro-aging phenotype (Shi & Murphy 2014), but the signal from the germline to the soma that conveys mating status is unknown.

**Figure 1.**
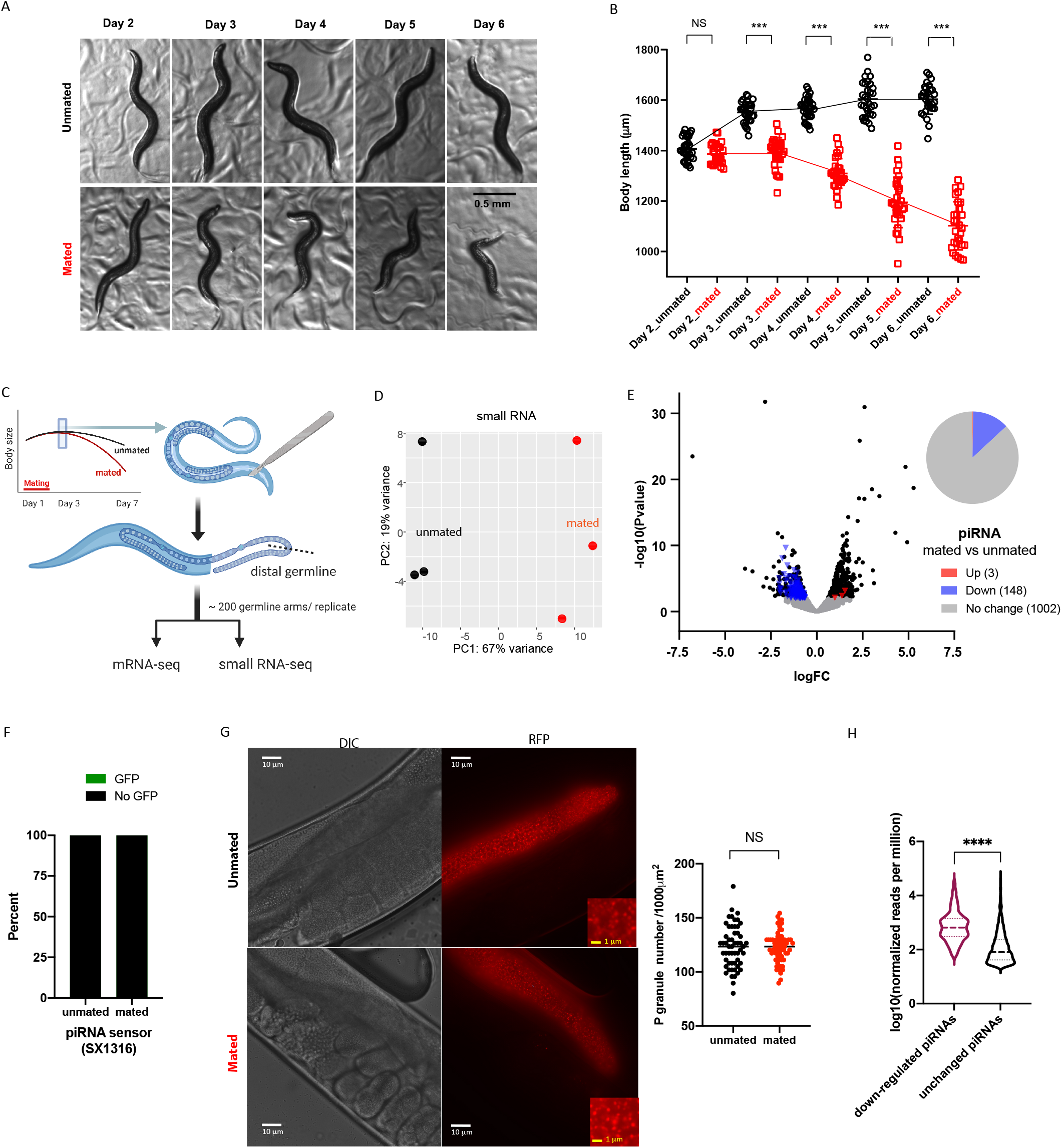
Mating induces shrinking and significant transcriptional changes in the germline. A) Representative pictures of the unmated and mated hermaphrodites from Day 2 – Day 6 of adulthood. Mating occurs on Day 1 for 24 hours. B) Length of unmated and mated fog-2 hermaphrodites. Two-tailed t-test was used for all body size measurement comparisons in this study, * p<0.05, ** p<0.01, *** p<0.001, NS: no statistical difference. C) Experimental design of germline isolation from mated and unmated fog-2 hermaphrodites. D) Principal Component Analysis (PCA) of normalized read counts from the small RNA transcriptomes of the isolated germlines E) Volcano plot of small RNA-seq transcriptome data of mated vs unmated germlines. Significantly differentially expressed piRNAs (P-adj ≤ 0.05) are highlighted in red (up-regulation) and blue (down-regulation). The pie chart (right) summarizes the number of significantly differentially expressed and unchanged piRNAs in the mated germline compared to the unmated germline. F) The piRNA sensor (SX1316) remains silenced in mated worms. 25 worms were checked for each condition. G) Mating does not affect the morphology and density of P granules. Left: representative images of mated and unmated Day 3 YY968 worms (P granules are labeled by PGL-1::RFP). Mating started on Day 1 for 2 days. Scale bar: 10 um (white), 1 um(yellow). Right: quantification of P granule density (blindly scored). H) piRNAs that are down-regulated after mating are initially expressed at a significantly higher level in the germline than piRNAs with no change.

To identify the germline pro-aging signal that induces post-mating physiological changes and early death, we dissected distal germlines from mated and unmated hermaphrodites, and extracted total RNA for both mRNA and small RNA sequencing (**Figure 1C**). Hermaphrodites were mated with young males from Day 1 of adulthood for two days before germline dissection. Differential expression analysis identified 666 significantly up-regulated and 590 down-regulated genes in the mated germline compared to the unmated germline (**Figure S1B-C**, Table 1). These genes were enriched in germline, reproductive system, and gonadal primordium function, confirming our successful dissection of the germline with minimum contamination by other tissues (**Figure S1D**). Genes up-regulated in the mated germline were enriched for ribosome, cuticle, protein hetero-dimerization, and peptide biosynthesis (**Figure S1E**), all of which are essential for rapid germ cell proliferation and cellularization. Genes down-regulated in the mated germline belonged to various metabolic categories such as rRNA metabolic process, cellular aromatic compound metabolic process, and heterocycle metabolic process (**Figure S1F**), indicating a shift of metabolic activities in the mated germline.

### piRNAs are significantly down-regulated in the mated germline

PIWI-associated RNAs (piRNAs) are predominantly expressed in the germline and are required to maintain germline integrity and fertility (Klattenhoff and Theurkauf, 2008). Previous studies have also shown that microRNAs are critical in lifespan regulation after germline removal (Boulias and Horvitz, 2012, Shen et al., 2012). Therefore, we performed small RNA sequencing from the same samples of dissected mated and unmated germlines. Differential expression analysis identified 440 significantly up-regulated and 303 down-regulated small RNAs in the mated germline compared to the unmated germline (**Figure 1D, S1H,** Table 2). Whereas miRNAs and other types of non-coding RNAs showed equal distribution in up- and down-regulation (**Figure S1G**), surprisingly, piRNAs were almost exclusively (98%) down-regulated in the mated germline (148 down-regulated vs 3 up-regulated; **Figure 1E, S1I**), indicating that mating triggers a unique response in germline piRNA abundance.

Mating does not appear to disrupt or inhibit the piRNA pathway, because mated and unmated worms showed no difference in piRNA sensor activity (the sensor is desilenced if the piRNA pathway is completely nonfunctional (Bagijn et al., 2012, Belicard et al., 2018); **Figure 1F**). Nor were P granule density and morphology affected by mating (P granules are essential for piRNA-mediated gene silencing (Ouyang et al., 2019); **Figure 1G**). Moreover, although we detected more than 1000 piRNAs in our dissected germlines that were unchanged upon mating, these 148 down-regulated piRNAs were initially expressed at a significantly higher level in the germline (**Figure 1H**), suggesting that mating leads to the down-regulation of a small and specific set of piRNAs without a general disruption of the piRNA pathway.

### Mating-induced shrinking and transcriptional changes require a functional piRNA pathway in the hermaphrodite germline

To further test which types of small RNAs are required for germline-specific mating-induced response, we measured body size changes after mating for different types of small RNA pathway mutants. Mating-induced shrinking requires germline hyperactivity and is a germline-specific aging phenotype (Shi and Murphy, 2014); therefore, shrinking serves as a good phenotypic readout for the germline-emanating pro-aging signal. We mated mutant hermaphrodites with young males for 2 days starting on Day 1 of adulthood and measured their body size on Day 6/7 of adulthood (**Figure S2A**). DCR-1 is required for both miRNA and siRNA processing (Grishok et al., 2001, Ketting et al., 2001, Knight and Bass, 2001). Like wild-type animals, after mating, *dcr-1* hermaphrodites still shrank (**Figure 2A, S2A**), suggesting that Dicer function is unnecessary for the germline-to-soma shrinking signal. Likewise, mutants of the dsRNA transporter SID-1 were also susceptible to mating-induced shrinking (**Figure 2A**), suggesting that mating-induced shrinking does not rely on functional miRNA and siRNA pathways.

**Figure 2.**
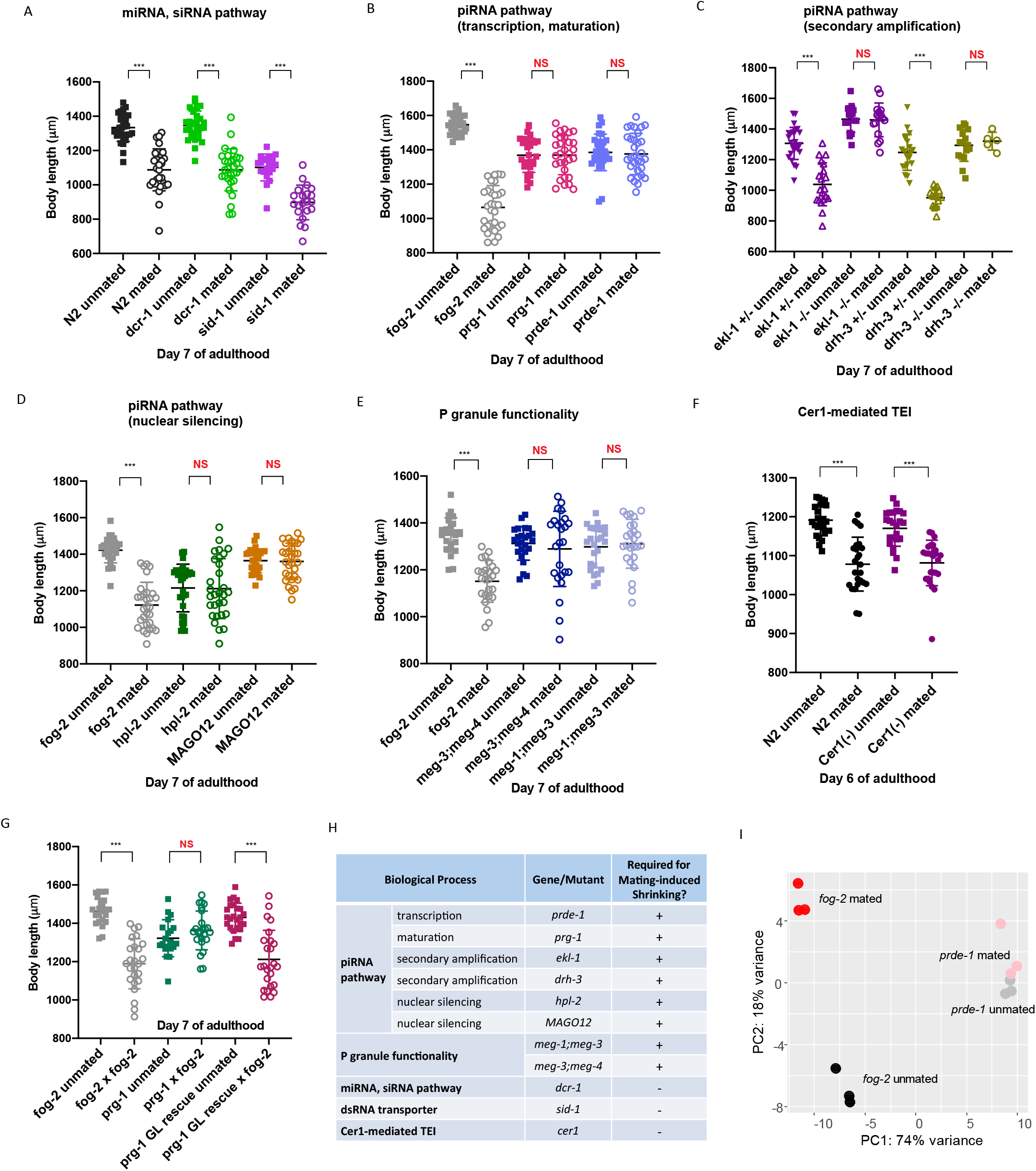
Mating-induced shrinking and transcriptional changes require the functional piRNA pathway in the germline. A-F) Body size measurements of mated and unmated hermaphrodites related to various types small RNA pathways. A) Mutants that are defective in miRNA or siRNA pathways still shrink after mating. N2 unmated: 1334±81 μm(n=30), N2 mated: 1088±122 μm(n=31), p<0.001. dcr(mg375) unmated: 1347±84 μm(n=30), mated: 1088±122 μm(n=30), p<0.001. sid-1(qt9) unmated: 1102±78 μm(n=20), mated: 898±101 μm(n=21), p<0.001. B-F) Mutants that affect multiple steps of the piRNA pathway are resistant to mating-induced shrinking. B) Ťog-2(q71) unmated: 1546±55 μm(n=30), mated: 1065±127 μm(n=30), p<0.001. prg-1(n4357) unmated: 1368±100 μ m(n=30), mated: 1369±114 μm(n=30), p=0.9603. prde-1(mj207) unmated: 1385±106 μm(n=30), mated: 1376±120 μm(n=31), p=0.7581. C) ekl-1 and drh-3 mutant alleles are maintained by chromosome balancers. Post-mat ing body size ofthe siblings with homozygous mutant alleles (-/-; no balancer) and heterozygous mutant allele (+/-, with the balancer) was measured in the same experiment. ekl-1(ok1197)[+/-] unmated: 1306±105 μm(n=25), mated: 1039±140 μm(n=20), p<0.001.ekl-1(ok1197)[-/-] unmated: 1464±87 μm(n=16), mated: 1460±111 μm(n=16), p=0.9063. drh-3(tm1217)[+/-] unmated: 1247±117μm(n=21), mated: 951±58 μm(n=16), p<0.001. drh-3(tm1217)[-/-] unmated: 1294±104 μm(n=16), mated: 1321±58 μm(n=5), p=0.5863. D) Ťog-2(q71) unmated: 1422±69 μm(n=30), mated: 1122±125 μm(n=30), p<0.001. hpl-2(ok916) unmated: 1215±130 μm(n=30), mated: 1212±164 μm(n=30), p=0.9286. MAGO12 unmated: 1365±65 μm(n=30), mated: 1361±100 μm(n=30), p=0.8577. E) Ťog-2(q71) unmated: 1349±73 μm(n=25), mated: 1151±89 μm(n=25), p<0.001. meg-3(tm4259) meg-4(ax2026) unmated: 1312±71 μm(n=25), mated: 1289±161 μm(n=25), p=0.5179. meg-1(vr10) meg-3(tm4259) unmated: 1298±89 μm(n=25), mated: 1311±104 μm(n=25), p=0.6330. F) N2 unmated: 1191±40 μm(n=25), mated: 1078±69 μm(n=25), p<0.001. Cerl(-)(RNAi for 3 generations) unmated: 1170±46 μm(n=25), mated: 1081±59 μm(n=25), p<0.001. G) Mating induces shrinking in hermaphrodites with germline-specific rescue of the piRNA pathway. Ťog-2(q71) unmated: 1462±68 μm(n=25), mated: 1189±131 μm(n=25), p<0.001. prg-1(n4357) unmated: 1322±96 μm(n=20), mated: 1363±101 μm(n=25), p=0.1803. Germline-specific prg-1 rescue (CQ655) unmated: 1431±74 μm(n=25), mated: 1212±152 μm(n=25), p<0.001. H) Summary of the genetic requirement for mating-induced shrinking. I) Principal Component Analysis (PCA) of normalized read counts from the mRNA transcriptomes of the isolated germlines. Black: fog-2 unmated, red: fog-2 mated, grey: prde-1 unmated, pink: prde-1 mated.

Next, we tested mutants for factors in multiple steps of the piRNA pathway, including piRNA transcription (*prde-1*), maturation (*prg-1*), secondary amplification (*ekl-1*, *drh-3*), and nuclear silencing (*hpl-2*) (Weick and Miska, 2014), as well as MAGO12, which has loss-of-function mutations in multiple Argonaute protein-encoding genes (Weick and Miska, 2014, Gu et al., 2009). Unlike the mutants involved in miRNA or siRNA processing (**Figure 2A**), none of these piRNA pathway mutants shrank after mating (**Figure 2B-D, S2A**), indicating that each gene in the piRNA pathway is required for mating-induced shrinking in the hermaphrodites. Moreover, one genomic copy of functional corresponding piRNA gene restored worms’ susceptibility to mating-induced shrinking (**Figure 2C**, *drh-3*, *ekl-1*, indicated by”+/-”). Together, our data suggest that the piRNA pathway function is critical for the signal that conveys the mated state of the germline to the soma.

piRNAs scan most of the transcriptome while located in P granules (*C. elegans* germ granules) (Shen et al., 2018, Zhang et al., 2018, Ouyang et al., 2019), and disruption of P granules compromises piRNA-mediated silencing (Ouyang et al., 2019). P granule distribution and assembly/disassembly dynamics depend on MEG proteins (Wang et al., 2014). We found that several *meg* double mutants in which P granule functionality is severely impaired (Wang et al., 2014) were also resistant to mating-induced shrinking (**Figure 2E**), demonstrating again that the germline-mediated postmating shrinking requires a fully functional piRNA pathway and P granules.

The *Cer1* retrotransposon was recently found to be required for communication between the germline and neurons in the transgenerational inheritance of pathogen avoidance, which also requires the germline and piRNAs (Moore et al., 2020). However, we found that worms lacking *Cer1* retrotransposon-encoded capsids still shrank after mating (**Figure 2F**), suggesting that *Cer1* is not involved in mating-induced shrinking.

Although piRNAs are predominantly expressed in the germline, there is emerging evidence that the piRNA pathway could also function in somatic tissue (Kim et al., 2018). To test whether the piRNA pathway is required autonomously in the germline to mediate mating-induced shrinking, we performed the same post-mating body size measurement using a germline-specific *prg-1* rescue strain, in which the only tissue with a functional piRNA pathway is the germline. We found that mating was able to induce shrinking in *prg-1* germline rescue strain (**Figure 2G**), confirming that piRNAs function in the germline to mediate post-mating shrinking (**Figure 2H**).

Next, we wondered whether the piRNA pathway is required for the dramatic mRNA transcriptional changes in mated germline. To address this question, we isolated mated and unmated distal germlines from *prde-1* mutants, which are defective in piRNA biosynthesis (Weick et al., 2014), and performed mRNA sequencing. Principal component analysis revealed that the lack of a functional piRNA pathway eliminated the transcriptional differences between the mated and unmated germline in *prde-1* worms (**Figure 2I**). The 660 significantly up-regulated genes previously identified in mated *fog-2* germlines were no longer up-regulated in *prde-1* mated germline (**Figure S2B**), indicating that the mating-induced transcriptional changes in the germline are also dependent on a functional piRNA pathway.

### Hedgehog signaling may encode the mysterious germline-originated pro-aging signal

Both post-mating shrinking and transcriptional changes require piRNA pathway function, implying that the germline pro-aging signal must be related to piRNAs. Since the pro-aging signal is amplified in mated germlines, while piRNAs are down-regulated, piRNAs themselves are unlikely to be the direct signal. One major role of piRNAs is gene silencing (Weick and Miska, 2014). Mating-induced down-regulation of piRNAs may release the silencing effect on piRNA targets, up-regulating the expression of these genes, making piRNAs’ targeted genes more likely candidates for the germline pro-aging signal.

To identify the signal, we took several steps to narrow the list of candidates (**Figure 3A**). Reasoning that the signal should be amplified after mating, we started with the list of 666 genes that are significantly up-regulated in the mated germlines we identified in our mRNA-seq analysis. First, since the pro-aging signal originates from the germline, it must be germline-specific: we subtracted those genes that are also up-regulated in mated *glp-1* germlineless worms (Booth et al., 2020) from the original list, reducing the number of candidates to 418 (**Figure S2C, 3A,** Table 3). Next, we reasoned that the amplified germline signal should be robust enough to be identified in the transcriptome of whole worms with an intact germline, since the germline accounts for a significant proportion of the whole worm biomass. We compared these 418 genes to our previous *fog-2* mated vs unmated whole worm genome-wide microarrays and found that the two datasets share 117 genes (Table 4). Furthermore, since piRNAs are required for mating-induced transcriptional changes and shrinking, the germline signal should also be dependent on piRNAs. Therefore, we cross-compared the remaining 117 genes with the potential targets (piRTarBase;(Wu et al., 2019)) of 148 significantly down-regulated piRNAs identified in the mated germline. We found that 34 of 117 genes were targets regulated by the 148 piRNAs that were down-regulated in mated germline (110 cl:2218 Table 5). Lastly, in order to relay the signal from the germline to the rest of the body, the gene product is likely to be secreted. We applied two prediction algorithms (Euk-mPLoc 2.0, SignalP) to the 34 genes and found that 13 are predicted to encode secreted proteins. We then ranked the final 13 genes according to their fold change of expression level in the germline, in the whole worm, and their consistency in expression patterns across the samples (**Figure 3B**, Table 5).

**Figure 3.**
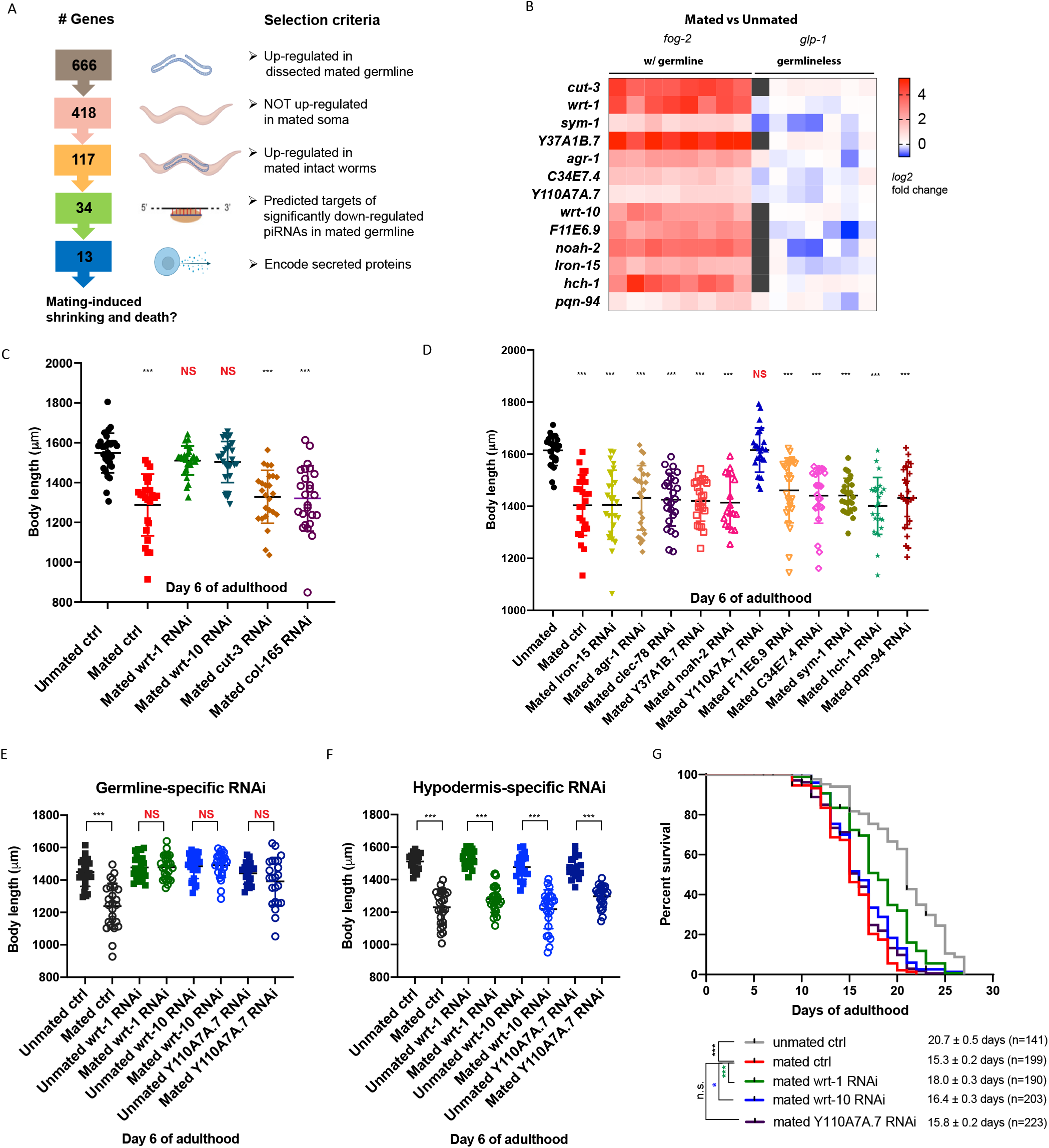
Secreted Hedgehog-like ligands in the germline are required for mating-induced shrinking and early death. A) Scheme describing the strategy and criteria to narrow the list of genes that might encode the germline pro-aging signal. B) Heatmap of the final 13 germline pro-aging signal candidates. Their mating-induced up-regulation is completely lost in mated germlineless worms. The data are displayed as log2 fold change of expression level in mated vs unmated whole worms. Left: fog-2 worms with functional germlines; right: glp-1 germlineless worms. C-D) Body size measurements of mated worms with RNAi treatment of individual candidate genes. C) fog-2(q71) unmated(control RNAi): 1549±99 μm(n=30); mated(control RNAi): 1288±155 μm(n=25), p<0.001;mated(wrt-1 RNAi): 1511±72 μm(n=27), p=0.1070; mated(wrt-10RNAi): 1503±103 μm(n=28), p=0.0917; mated(cut-3 RNAi): 1329±133 μm(n=25), p<0.001; mated(col-165 RNAi): 1321±165 μm(n=25), p<0.001. D) Ťog-2(q71) unmated(control RNAi): 1615±58 μm(n=25); mated(control RNAi): 1404±116 μm(n=25), p<0.001; mated(lron-15 RNAi): 1405±133 μm(n=25), p<0.001; mated(agr-1 RNAi): 1433±124 μm(n=25), p<0.001;mated(clec-78 RNAi): 1425±101 μm(n=25), p<0.001;mated(Y37A1B.7 RNAi): 1421±78 μm(n=25), p<0.001; mated(noah-2 RNAi): 1414±100 μm(n=25), p<0.001; mated(Y110A7A.7 RNAi): 1615±85 μm(n=25), p=0.9780; mated(F11E6.9 RNAi): 1461±124 μm(n=25), p<0.001;mated(C34E7.4RNAi): 1441±106 μm(n=25), p<0.001;mated(sym-1 RNAi): 1441±66 μm(n=25), p<0.001; mated(hch-1 RNAi): 1401±109 μm(n=25), p<0.001; mated(pqn-94 RNAi): 1433±118 μm(n=25), p<0.001. E-F) Germline-specific knock down (E) but not hypodermis-specific knock down (F) of wrt-1, wrt-10, and Y110A7A.7 protectsthe mated hermaphroditesfrom shrinking. E) Germline-specific RNAi strain(DCL569) unmated(control RNAi): 1449±89 μm(n=25), mated(control RNAi): 1239±141 μm(n=25), p<0.001; unmated(wrt-1 RNAi): 1477±71 μm(n=25), mated(wrt-1 RNAi): 1479±74 μm(n=25), p=0.9211;unmated(wrt-10 RNAi): 1485±76 μm(n=25), mated(wrt-10 RNAi): 1491±77 μm(n=25), p=0.7701;unmated(Y110A7A.7 RNAi): 1441±66 μm(n=20), mated(Y110A7A.7 RNAi): 1391±147 μm(n=23), p=0.1698. F) Hypodermis-specific RNAi strain(CQ479) unmated(control RNAi): 1511±51 μm(n=22), mated(control RNAi): 1230±105 μm(n=26), p<0.001; unmated(wrt-1 RNAi): 1528±49 μm(n=25), mated(wrt-1 RNAi): 1282±83 μm(n=25), p<0.001; unmated(wrt-10 RNAi): 1477±75 μm(n=21), mated(wrt-10 RNAi): 1218±120 μm(n=26), p<0.001; unmated(Y110A7A.7 RNAi): 1474±67 μm(n=17), mated(Y110A7A.7 RNAi): 1297±67 μm(n=24), p<0.001. G) RNAi knockdown of wrt-1 and wrt-10 increasesthe lifespan of mated worms, (more replicatesareshown in Figure S3) For all the lifespan assays performed in this study, Kaplan-Meier analysis with log-rank (Mantel-Cox) test was used to determine statistical significance. Mated control: 15.3±0.2 days, n=199; unmated control: 20.7±0.5days, n=141, p<0.001; mated wrt-1 RNAi: 18.0±0.3days, n=190, p<0.001; mated wrt-10 RNAi: 16.4±0.3days, n=203, p=0.0214; mated Y110A7A.7 RNAi: 15.8±0.2days, n=223, p=0.0709. All lifespans were compared to that of mated worms treated with control (L4440 vector) RNAi, *p < 0.05, ***p < 0.001, n.s., not significant.

To determine which of these 13 genes might be the germline pro-aging signal, we measured body size postmating after RNAi treatment for each individual gene. Knockdown of three candidates prevented the mated worms from shrinking: two warthog (Hedgehog-like family) genes, *wrt-1*, *wrt-10*, and an uncharacterized gene, Y110A7A.7 (**Figure 3C, D**). Knocking down these three genes specifically in the germline (**Figure 3E**) but not in the hypodermis (**Figure 3F**) was sufficient to protect the mated hermaphrodites from shrinking, confirming that they are specifically required in the germline to mediate mating-induced shrinking.

In addition to preventing post-mating shrinking, inhibiting the *bona fide* germline pro-aging signal should also increase the lifespan of mated worms. Y110A7A.7 RNAi did not prevent the worms from mating-induced death (**Figure 3G, S3A-B**). By contrast, reduction of *wrt-1* provided over 50% protection against mating-induced early death, and knocking down *wrt-10* yielded a 20% protection (**Figure 3G, S3C-D**, **4D**). Similarly, mating-induced death was largely attenuated in piRNA pathwaydeficient mutants in which the germline pro-aging signal is abrogated (**Figure S3E-F**). The remaining unrescued lifespan decrease is likely caused by the germlineindependent mating-induced death mechanisms downstream of seminal fluid transfer (Shi and Murphy, 2014, 2021). Our results suggest that the piRNA pathway and its downstream Hegdehog signaling specifically encode the germline pro-aging signal.

To identify possible receptors for the Hedgehog ligands secreted from the germline upon mating, we tested the roles of Patched receptor homolog genes in shrinking after mating. Knocking down *ptr-6*, *ptr-10*, and *ptr-16* consistently prevented mated worms from shrinking (**Figure 4A**), making them the likely candidates to receive the germline-originating signal encoded by *wrt-1* and *wrt-10*. Most hedgehog signaling receptors, including *ptr-6*, *ptr-10*, and *ptr-16*, are predicted to be expressed in the hypodermis (**Figure 4B, S4A** (Chikina et al., 2009, Yao et al., 2018)), suggesting that hypodermis is the major receiving tissue of the germline pro-aging signal mediating post-mating shrinking and death. Knocking down *ptr-6* and *ptr-16*, but not *ptr-10* or the other Patched-related receptor genes, significantly rescued mating-induced early death (**Figure 4C, S4B, F-O**). *Ptr-6* and *ptr-16* RNAi yielded a similar or greater degree of protection compared to *wrt-1* RNAi in mated worms (**Figure 4D**), suggesting that they are likely to be the receptors that receive the pro-aging Hedgehog signal from the germline.

**Figure 4.**
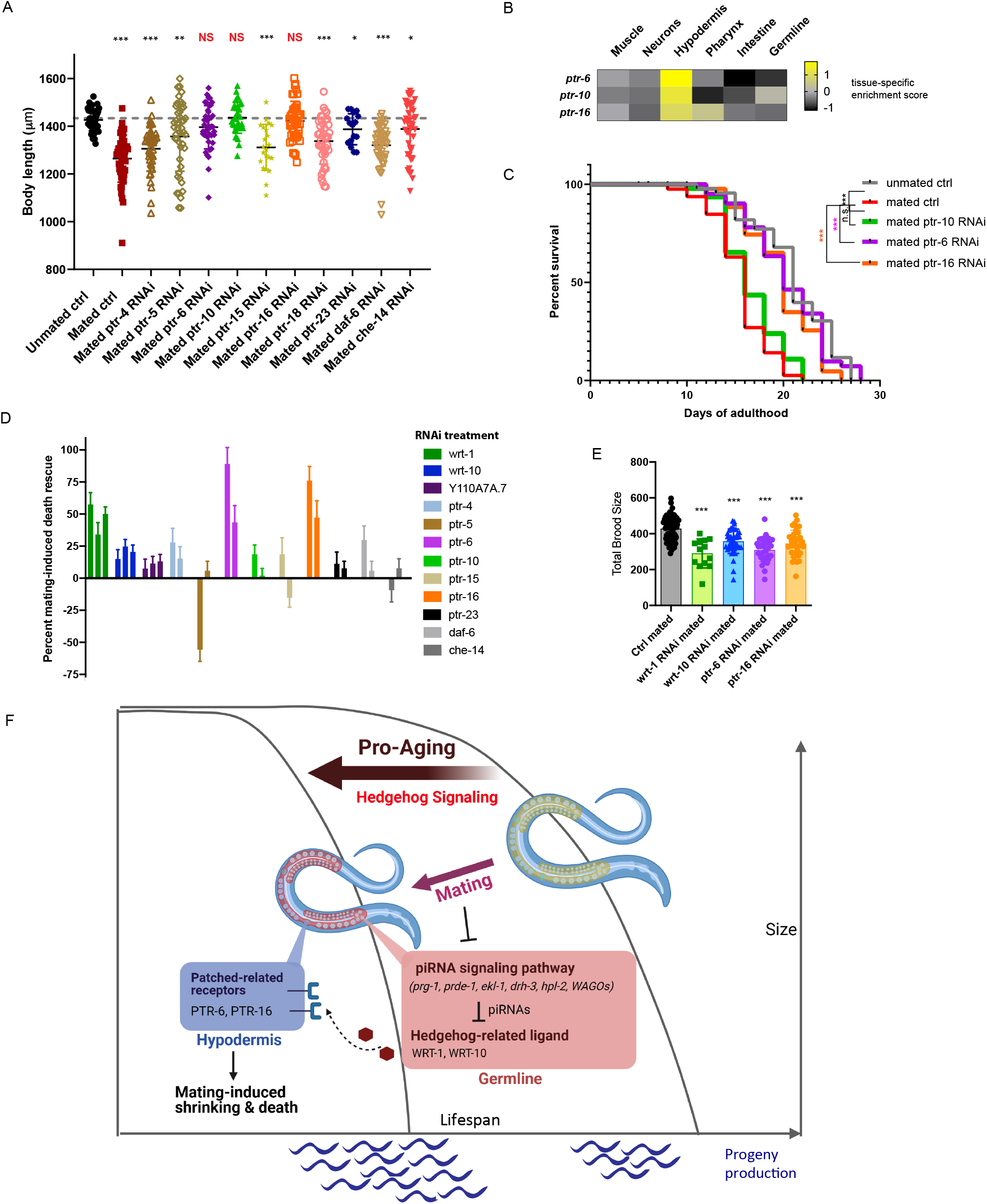
Hedgehog signaling encodes the germline-to-soma aging signal in mated hermaphrodites. A) Body size measurements of mated worms with RNAi treatment of Patched receptor homolog genes, fog-2(q71) unmated(control RNAi): 1426±46 μm(n=44); mated(control RNAi): 1264±99 μm(n=48), p<0.001;mated(ptr-4 RNAi): 1305±94 μm(n=43), p<0.001;mated(ptr-5 RNAi): 1357±146 μm(n=45), p=0.0032; mated(ptr-6 RNAi): 1396±91 μm(n=42), p=0.0502; mated(ptr-10 RNAi): 1436±65 μm(n=34), p=0.4623; mated(ptr-15 RNAi): 1311±96 μm(n=22), p<0.001; mated(ptr-16 RNAi): 1422±83 μm(n=43), p=0.7409; mated(ptr-18 RNAi): 1337±96 μm(n=42), p<0.001; mated(ptr-23 RNAi): 1387±65 μm(n=17), p=0.0104; mated(daf-6 RNAi): 1320±86 μm(n=44), p<0.001; mated(che-14 RNAi): 1388±112 μm(n=44), p=0.0394. B) Tissue-specific expression prediction of ptr-6, ptr-10, and ptr-16. C) Knock down of ptr-6 and ptr-16, but not ptMOrescues the mating-induced death of fog-2 hermaphrodites. Mated control: 15.7±0.3 days, n=85; unmated control: 21.1±0.6 days, n=46, p<0.001; mated ptr-10 RNAi: 1θ.7±0.4 days, n=50, p=0.0807; mated ptr-6 RNAi: 20.5±0.7 days, n=46, p<0.001; mated ptr-16 RNAi: 19.8±0.6 days, n=50, p<0.001. All lifespans were compared to that of mated worms treated with control RNAi, ***p < 0.001, n.s., not significant. D) Summary of mating-induced death rescue effect of RNAi knock down of individual Hedgehog signaling components. Each bar of the same color represents one biological replicate of the lifespan assay. E) Inhibiting the Hedgehog signaling by RNAi in mated N2 hermaphrodites leads to significantly reduced total mated brood size. Control mated: 429±67, n=65; wrt-1 RNAi mated: 290±8I, n=13; wrt-10 RNAi mated: 358±68, n=41; ptr-6 RNAi mated: 3IO±65, n=33; ptr-16 RNAi mated: 345±76, n=38. ***, p<0.001, t-test was used for total mated brood size comparison (all were compared to that of control mated worms). F) Model of piRNA-mediated Hedgehog signaling as the germline-to-soma pro-aging signal in mated hermaphrodites

Inhibiting germline piRNA-mediated Hedgehog signaling is beneficial to the mothers, as it prevents mated worms from shrinking and largely rescues mating-induced death. However, such benefits do not come without a cost: suppression of the germline pro-aging signal leads to a reduced brood size. The mated brood size of *prde-1* mutants was less than 20% of wild-type’s (**Figure S4C**). RNAi knockdown of the components of Hedgehog signaling in mated wild-type worms also led to a 20%-30% reduction in brood size (**Figure 4E**; p<0.001)

## Discussion

In animals across taxa, germline hyperactivity leads to accelerated aging (Shi and Murphy, 2021), however, the underlying mechanisms are poorly understood. Here we have shown that mating-induced piRNA down-regulation in the germline releases the suppression of the Hedgehog signaling pathway, which in turn leads to body size and lifespan decrease in mated worms (**Figure 4F**). Our results suggest that piRNA-regulated Hedgehog signaling encodes the previously unknown germline-to-soma pro-aging signal (Hsin and Kenyon, 1999).

The Hedgehog signaling pathway is one of the key regulators of animal development (Ingham et al., 2011); our study provides evidence for an additional critical role in adult post-mating lifespan regulation in *C. elegans*. In addition to its role in the germline-to-soma signaling we report here, Hedgehog signaling also participates in soma-to-germline communication: overexpression of hypodermal *wrt-10* is sufficient to delay reproductive decline and improve germline health (Templeman et al., 2020). Therefore, *wrt-10* could function as a feedforward loop in mated worms, achieving sustained germline hyperactivity at the cost of exacerbated somatic collapse (i.e. shrinking). Our results suggest that there is likely a trade-off between somatic integrity and progeny production in mated animals.

The involvement of Hedgehog signaling in lifespan regulation is not limited to worms. Recently, it has been reported that both enhanced and reduced Hedgehog signaling lead to reduced survival of *Drosophila* larvae under starvation (Rodenfels et al., 2014). Impaired Hedgehog signaling in glia but not in neurons affects the lifespan of adult Drosophila (Rallis et al., 2020). Moreover, the Hedgehog pathway inhibitor saridegib dramatically increases lifespan by four-fold in a mouse medulloblastoma model (Lee et al., 2012). Therefore, Hedgehog signaling seems to have an evolutionarily conserved role in lifespan regulation beyond its well-established function in development.

Our results unveil a new function for the piRNA pathway in transcriptional regulation, an underappreciated aspect of piRNA biology compared to its better-studied role in repression of transposable elements (TEs) in animal germlines (Ozata et al., 2019). Previous studies regarding piRNAs’ TE-independent functions in endogenous mRNA regulation have uncovered roles in various developmental processes including embryonic patterning, germ cell specification, and stem cell biology (Rojas-Ríos and Simonelig, 2018). Our results expand the role of the piRNA pathways in the regulation of gene expression to an adult function, which is largely unstudied except in neurons (Lee et al., 2011, Rajasethupathy et al., 2012, Kim et al., 2018). Our results provide the first observation of piRNA-mediated regulation of a known signaling pathway in the adult germline. These results also highlight the fact that piRNA expression is regulated, tunable, and can respond to changes in mating status. The cell-non-autonomous nature of downstream Hedgehog signaling extends the influence of piRNAs beyond the adult germline to somatic tissues, indicating a mechanism the piRNA pathway employs to regulate biological processes in other tissues that may lack piRNA machinery.

Mating-induced germline-to-soma piRNA-mediated Hedgehog signaling elegantly coordinates germline function and somatic aging. In an unmated female with low or no germline proliferation, the priority of the germline is to maintain its integrity until mating occurs, while avoiding unnecessary proliferation or differentiation of the germ cells. piRNAs contribute to this process by suppressing developmental programs, including those regulated by Hedgehog signaling. By contrast, upon mating, the germline switches to progeny production mode, and the previous suppression of Hedgehog signaling by piRNAs is released to allow rapid germline stem cell proliferation and further differentiation. As Hedgehog ligands are secreted, activated Hedgehog signaling may be co-opted as a germline-to-soma signal to coordinate the somatic responses to elevated germline activity. Post-mating shrinking correlates with the reallocation of somatic resources to the germline (Shi and Murphy, 2014, Shi et al., 2017, Heimbucher et al., 2020) and the reduction in energy devoted to somatic integrity maintenance, reflecting the direct cost of germline hyperactivity and significantly increased progeny production. Together, our study reveals a mechanism that can efficiently convey germline status to the soma in adulthood, allowing the animals to better organize the balance between reproduction and somatic maintenance, optimizing reproductive fitness.

## Acknowledgments

We thank the *Caenorhabditis* Genetics Center (CGC) for strains; the Genomics Core Facility at Princeton University; BioRender for model figure design software; and members of the CTM’s laboratory for critically reading the manuscript and valuable feedback. C.T.M. is the Director of the Glenn Center for Aging Research at Princeton and an HHMI-Simons Faculty Scholar. This work was supported by a Pioneer Award to C.T.M. (NIGMS DP1GM119167).

## Materials and Methods

### Strains

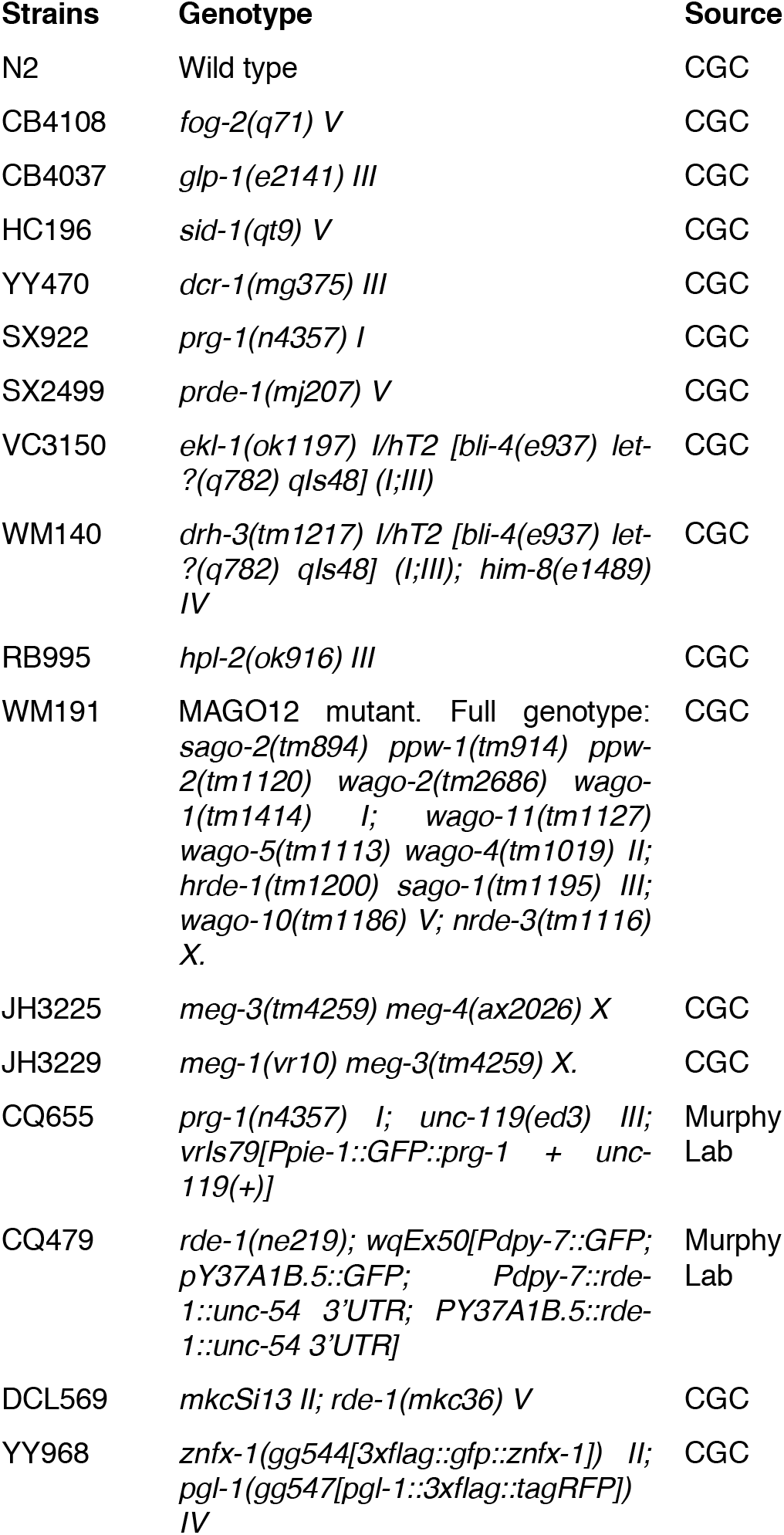

| Strains | Genotype | Source |
| --- | --- | --- |
| N2 | Wild type | CGC |
| CB4108 | *fog-2(q71) V* | CGC |
| CB4037 | *glp-1(e2141) III* | CGC |
| HC196 | *sid-1(qt9) V* | CGC |
| YY470 | *dcr-1(mg375) III* | CGC |
| SX922 | *prg-1(n4357) I* | CGC |
| SX2499 | *prde-1(mj207) V* | CGC |
| VC3150 | *ekl-1(ok1197) I/hT2 [bli-4(e937) let-?(q782) qls48] (I;III)* | CGC |
| WM140 | *drh-3(tm1217) I/hT2 [bli-4(e937) let-?(q782) qls48] (I;III); him-8(e1489) IV* | CGC |
| RB995 | *hpl-2(ok916) III* | CGC |
| WM191 | MAGO12 mutant. Full genotype: *sago-2(tm894) ppw-1(tm914) ppw-2(tm1120) wago-2(tm2686) wago-1(tm1414) I; wago-11(tm1127) wago-5(tm1113) wago-4(tm1019) II; hrde-1(tm1200) sago-1(tm1195) III; wago-10(tm1186) V; nrde-3(tm1116) X.* | CGC |
| JH3225 | *meg-3(tm4259) meg-4(ax2026) X* | CGC |
| JH3229 | *meg-1(vr10) meg-3(tm4259) X.* | CGC |
| CQ655 | *prg-1(n4357) I; unc-119(ed3) III; vrls79[Ppie-1::GFP::prg-1 + unc-119(+)]* | Murphy Lab |
| CQ479 | *rde-1(ne219); wqEx50[Pdpy-7::GFP; pY37A1B.5::GFP; Pdpy-7::rde-1::unc-54 3'UTR; PY37A1B.5::rde-1::unc-54 3'UTR]* | Murphy Lab |
| DCL569 | *mkcSi13 II; rde-1(mkc36) V* | CGC |
| YY968 | *znfx-1(gg544[3xflag::gfp::znfx-1]) II; pgl-1(gg547[pgl-1::3xflag::tagRFP]) IV* | CGC |

### Germline dissection

Germline dissection method was modified from the previous publication (Campbell and Updike, 2015). Worms were transferred to iced M9 buffer for dissection. Heads or tails were removed with 26G needles, allowing the distal portion of the germline to pop out of the worm. Glass capillaries pulled with an opening just large enough to fit the end of the germline were used to rapidly detach them at the ventral to distal bend. Dissected distal germlines were transferred immediately into 1.5 ml eppendorf tubes filled with 500 ul Trizol. About 200 germlines were collected for each biological replicate.

### RNA isolation and library preparation

Tubes containing dissected germlines (immersed in Trizol) were put in the Eppendorf MixMate Vortex Mixer at 800 rpm and 65 C for 1 hour before the isolation process. Total RNA was extracted from Trizol using the mirVana miRNA isolation kit (ThermoFisher). mRNA libraries for directional RNA sequencing were prepared using the SMARTer Apollo System and were sequenced (150-nt single-end) on the Illumina NovaSeq platform. RNA samples for small RNA-seq were treated with 5’polyphosphatase (Lucigen) and prepared using the SMARTer Apollo system with modifications for small RNA library preparation. Small RNA-containing libraries were Blue Pippin size selected (15-30nt insert size) prior to 75-nt single-end sequencing.

### Transcriptome data analysis

#### RNA-seq

FASTQC was used to assess read quality scores. Universal adaptor sequences were trimmed from small RNA library sequences using Cutadapt. Reads were mapped to the C. elegans genome (UCSC Feb 2013, ce11/ws245) using Bowtie. Count matrices were generated using featureCounts. Data were normalized with a variance-stabilizing transformation (DESeq2). DESeq2 was used for differential expression analysis. Genes at p-adj < 0.05 were considered significantly differentially expressed. Principal Component Analysis (PCA) was carried out using the R method (prcomp). Heatmaps were generated in R using normalized read counts (variance-stabilizing transformation). Tissue enrichment and GO term enrichment analysis were performed using wormbase enrichment analysis (https://wormbase.org/tools/enrichment/tea/tea.cgi) for significantly differentially expressed genes. Predicted targets of piRNAs were retrieved from piRTarBase (http://cosbi6.ee.ncku.edu.tw/piRTarBase/), see Table 5 for the full list. Sequences will be deposited at NCBI BioProject PRJNA752368.

#### Microarrays

Microarray data of mated and unmated *glp-1*(*e2141*) was retrieved from the previous publication (Booth et al., 2020). Microarray data of mated and unmated *fog-2*(*q71*) was retrieved from the previous publication (Choi et al., 2021). Hermaphrodites were mated on day 1 of adulthood for 24 h in a 2:1male:hermaphrodite ratio. About 200 hermaphrodites were collected on day 2/3 of adulthood for each biological replicate. RNA was extracted by the heat vortexing method. Two-color Agilent microarrays were used for expression analysis. Significantly differentially expressed gene sets were identified using SAM (Tusher et al., 2001). One class SAM was performed to identify genes that are significantly differentially expressed after mating. The lists were then compared to significantly up-regulated genes in dissected germline (RNA-seq).

### Lifespan

60 mm NGM plates were used to set up group mating. Each 60 mm NGM plate was seeded with OP50 to make a ~ 3 cm diameter bacterial lawn 2 days before mating. All lifespan assays were performed at room temperature (~20–21°C). About 50 hermaphrodites and 100 young (day 1 – day 2 of adulthood) males were transferred onto the plate. 24/48 hr later, the hermaphrodites were transferred onto newly seeded 60 mm NGM plates in the absence of males for lifespan assays. No FUdR was added to the plates. About 25 synchronized Day 1 hermaphrodites were transferred onto each plate. The hermaphrodites were transferred daily onto new seeded plates in the first week of the lifespan assay. Afterward, they were transferred once every 2 days. When RNAi was used in lifespan assay, RNAi treatment always started from eggs for all the experiments in this study. Kaplan-Meier analysis with log–rank (Mantel-Cox) method was performed to compare the lifespans of different groups. ‘Bagged’ worms were censored on the day of the event.

### Body size measurements

Images of live hermaphrodites on 60 mm plates were taken on Day 6/7 of adulthood with a Nikon SMZ1500 microscope. When RNAi was used, RNAi treatment always started from eggs for all the experiments. Image J was used to analyze the body size of the worms. The middle line of each worm was delineated using the segmented line tool and the total length was documented as the body length of the worm. T-test was performed to compare the body size differences between groups of worms in the same day.

### Brood size

Individual hermaphrodites after mating were transferred onto 3 cm NGM plates seeded with 25 mL of OP50 and moved to fresh plates daily until reproduction ceased. The old plates were left at 20C for 2 days to allow the offspring to grow into adults, which were manually counted for daily production and total brood size. 10-25 plates of individual worms of each genotype/treatment were counted to account for individual variation.

### P granules imaging and quantification

mCherry-tagged fluorescent PGL-1 were visualized in living nematodes (YY968) by mounting young adult animals on 2% agarose pads with M9 buffer with 20 mM levamisole, Fluorescent images were captured using a Nikon Ti microscope with a 100× objective. Images were processed and quantified in ImageJ. The quantification of germline granule fluorescence was performed using ImageJ. For every image, a region of interest (ROI) with a clear focus of P granules was selected manually. The area of the whole ROI was kept the same for all the images. The number of puncta within the ROI was measured blindly for each germline and image. The densities of germline P granules were calculated as: the number of puncta within the ROI/the area of the whole ROI. Densities of germline granules were determined for 50–60 ROIs, and the mean and SD were calculated using GraphPad Prism. T-test was performed to compare the P granule density differences between mated and unmated germlines.

## Supplementary Figures

**Figure S1.**
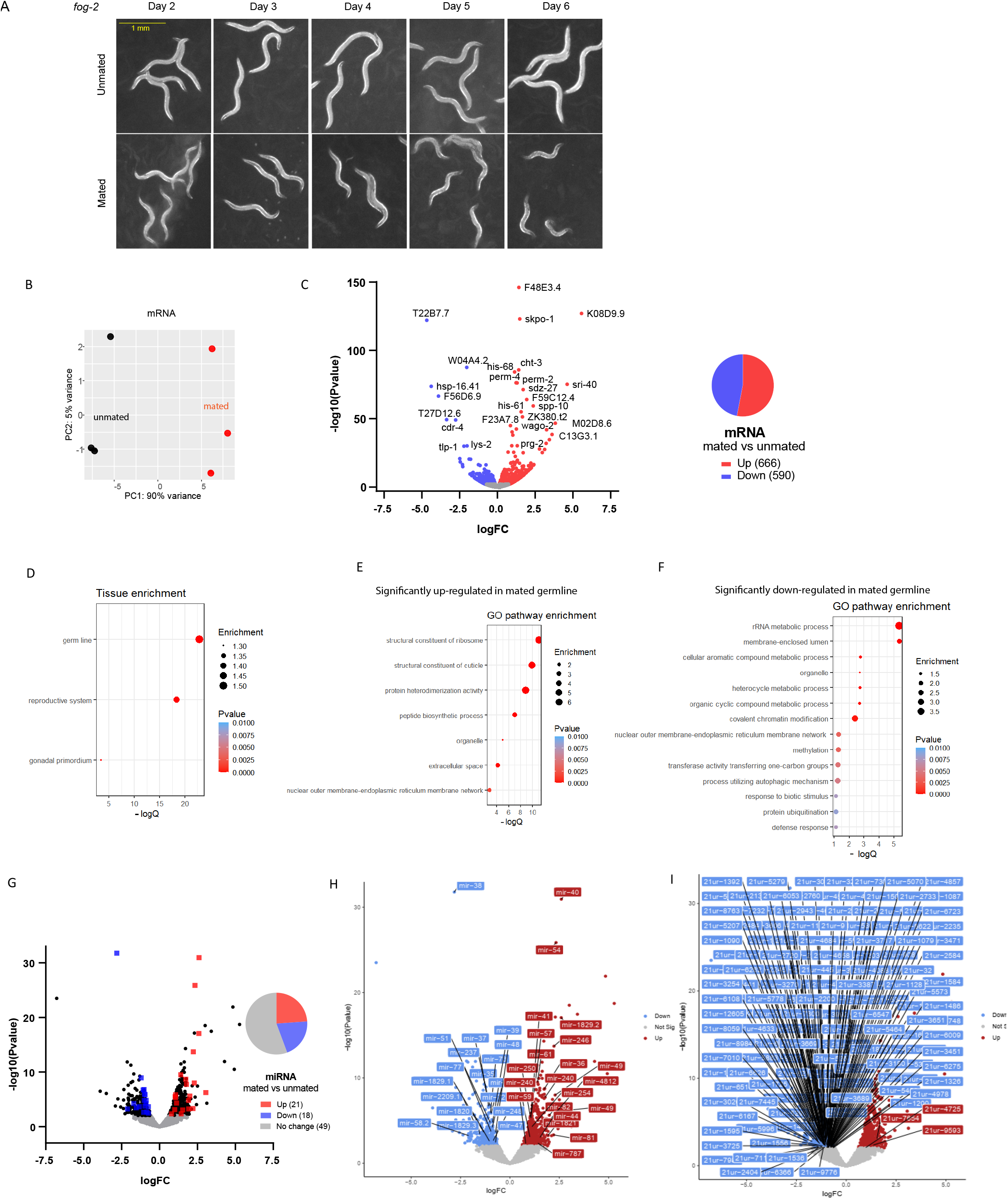
Mating-induced shrinking and analyses of mating-induced germline transcriptional changes. A) Representative pictures of the unmated and mated hermaphrodites from Day 2 – Day 6 of adulthood. Mating occurs on Day 1 for 24 hours. B) Principal Component Analysis (PCA) of normalized read counts from the mRNA transcriptomes of the isolated germlines. C) Volcano plot of mRNA-seq transcriptome data displaying the pattern of gene expression values for mated vs unmated germlines. Significantly differentially expressed genes (P-adj ≤ 0.05) are highlighted in red (up-regulation) and blue (down-regulation). The pie chart (right) summarizes the number of significantly differentially expressed genes in the mated germline compared to the unmated germline. D) Tissue enrichment analysis of significantly differentially expressed genes in the mated germline E-F) Gene Ontology (GO) enrichment analysis of significantly up- (E) and down- (F) regulated genes in the mated germline. (G) Volcano plot of small RNA-seq transcriptome data of mated vs unmated germlines. Significantly differentially expressed miRNAs (P-adj ≤ 0.05) are highlighted in red (upregulation) and blue (down-regulation). The pie chart (right) summarizes the number of significantly differentially expressed and unchanged miRNAs in the mated germline compared to the unmated germline. H-I) Volcano plot of small RNA-seq transcriptome data of mated vs unmated germlines. Significantly differentially expressed small RNAs (P-adj ≤ 0.05) are highlighted in red (upregulation) and blue (down-regulation). The top 15 significantly differentially expressed small RNAs (all types) in the mated germline are shown in H. I represents all significantly differentially piRNAs.

**Figure S2.**
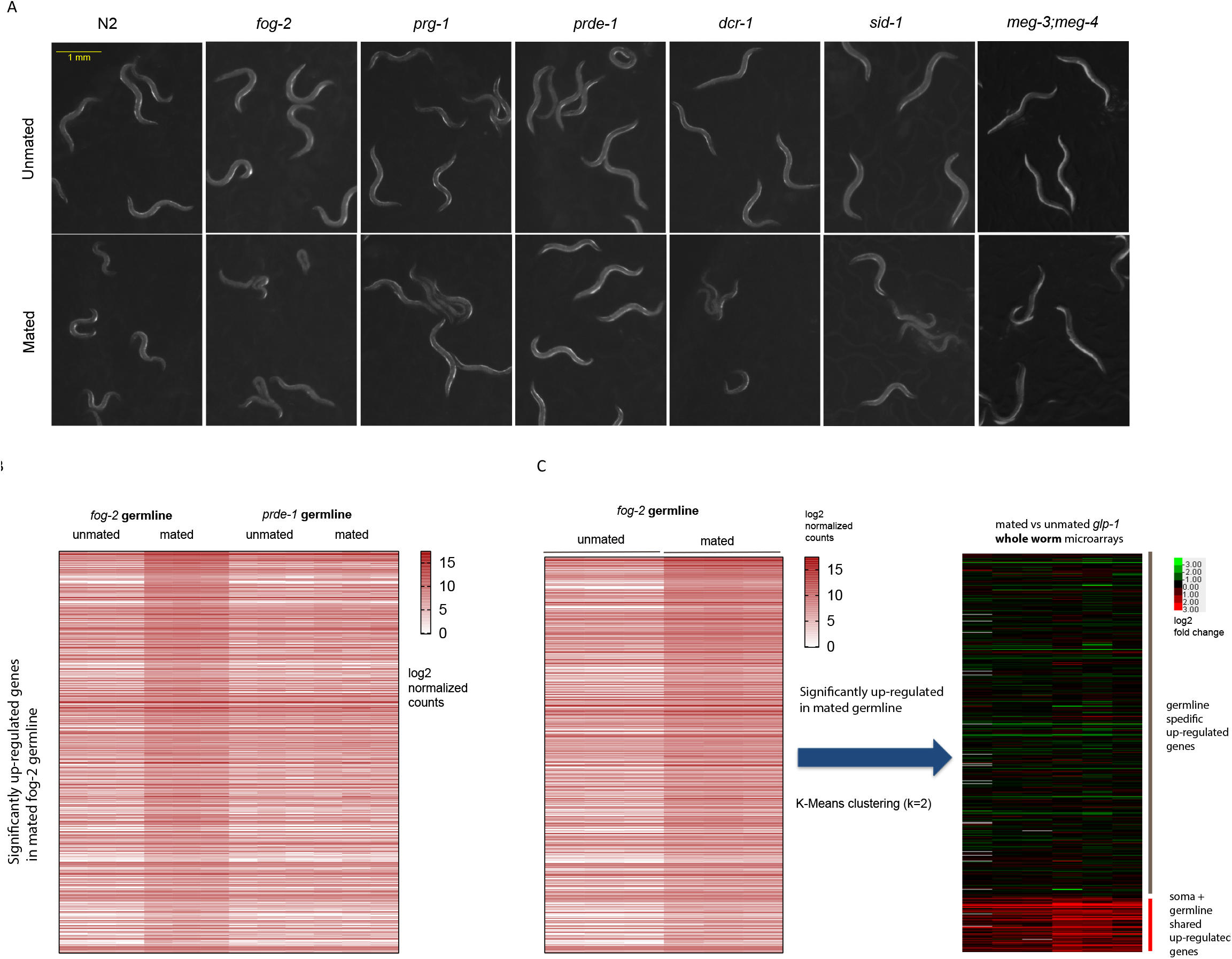
piRNA pathway mutants are resistant to mating-induced shrinking and lack transcriptional responses in the germline. A) Representative pictures of the unmated and mated hermaphrodites related to various types of small RNA pathways on Day 6/7 of adulthood. B) Heatmap of genes that are significantly up-regulated in mated *fog-2* germline. The data are displayed as log2(normalized counts). Such mating-induced up-regulation in expression is completely abolished in piRNA pathway-deficient *prde-1* germline. C) K-Means (k=2) clustering separates the significantly up-regulated genes in *fog-2* mated germline to germline-specific transcriptional response and generic transcriptional response to mating

**Figure S3.**
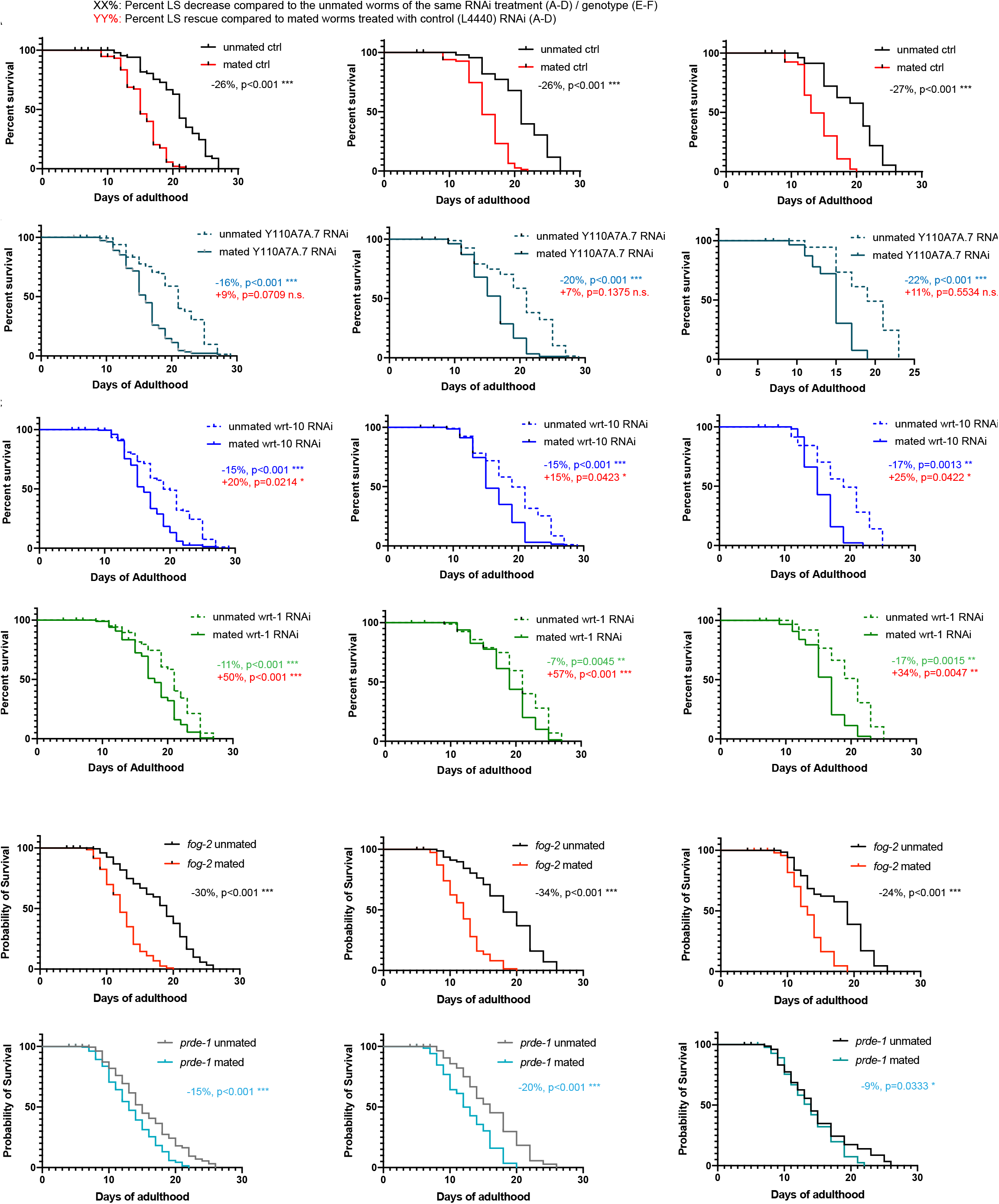
Inhibiting piRNA pathway and Hedgehog signaling largely ameliorates mating-induced early death. A-D) Lifespans of mated and unmated *fog-2*(*q71*) worms treated with control(A), Y110A7A.7(B), *wrt-10*(*C*), and *wrt-1*(D) RNAi. Replicates were shown in left, middle, and right panels. Kaplan–Meier analysis with log-rank (Mantel–Cox) test was used to determine statistical significance. A) Left, unmated ctrl: 20.7±0.5 days, n=141, mated ctrl: 15.3±0.2 days, n=199, p<0.001; Middle, unmated ctrl: 21.1±0.6 days, n=46, mated ctrl: 15.7±0.3 days, n=85, p<0.001; Right, unmated ctrl: 19.5±0.9 days, n=53, mated ctrl: 14.2±0.4 days, n=71, p<0.001. B) Left, unmated Y110A7A.7 RNAi: 18.8±0.5 days, n=196, mated Y110A7A.7 RNAi: 15.8±0.2 days, n=233, p<0.001(compared to unmated Y110A7A.7 RNAi), p=0.0709(compared to mated control); Middle, unmated Y110A7A.7 RNAi: 20.1±0.6 days, n=81, mated Y110A7A.7 RNAi: 16.1±0.4 days, n=105, p<0.001(compared to unmated Y110A7A.7 RNAi), p=0.1375(compared to mated control); Right, unmated Y110A7A.7 RNAi: 18.9±1.1 days, n=60, mated Y110A7A.7 RNAi: 14.8±0.3 days, n=61, p<0.001(compared to unmated Y110A7A.7 RNAi), p=0.5534(compared to mated control). C) Left, unmated wrt-10 RNAi: 19.3±0.4 days, n=237, mated wrt-10 RNAi: 16.4±0.3 days, n=203, p<0.001(compared to unmated wrt-10 RNAi), p=0.0214(compared to mated control); Middle, unmated wrt-10 RNAi: 19.3±0.5 days, n=108, mated wrt-10 RNAi: 16.5±0.4 days, n=80, p<0.001(compared to unmated wrt-10 RNAi), p=0.0423(compared to mated control); Right, unmated wrt-10 RNAi: 18.8±1.1 days, n=65, mated wrt-10 RNAi: 15.5±0.3 days, n=72, p<0.001(compared to unmated wrt-10 RNAi), p=0.0422(compared to mated control). D) Left, unmated wrt-1 RNAi: 20.2±0.4 days, n=180, mated wrt-1 RNAi: 18.0±0.3 days, n=190, p<0.001(compared to unmated wrt-1 RNAi), p<0.001(compared to mated control); Middle, unmated wrt-1 RNAi: 20.3±0.5 days, n=83, mated wrt-1 RNAi: 18.8±0.5 days, n=90, p=0.0045(compared to unmated wrt-1 RNAi), p<0.001(compared to mated control); Right, unmated wrt-1 RNAi: 19.4±0.8 days, n=60, mated wrt-1 RNAi: 16.0±0.5 days, n=63, p=0.0015(compared to unmated Y110A7A.7 RNAi), p=0.0047(compared to mated control). E-F) Lifespans of mated and unmated worms with defective piRNA pathway (F) and control (E). E) Left, *fog-2* unmated ctrl: 17.8±0.4 days, n=169, *fog-2* mated ctrl: 12.4±0.3 days, n=167, p<0.001; Middle, *fog-2* unmated ctrl: 18.2±0.6 days, n=85, *fog-2* mated ctrl: 12.0±0.3 days, n=83, p<0.001; Right, *fog-2* unmated ctrl: 17.4±0.6 days, n=84, *fog-2* mated ctrl: 13.2±0.4 days, n=84, p<0.001. F) Left, *prde-1* unmated ctrl: 15.7±0.4 days, n=164, *prde-1* mated ctrl: 13.4±0.3 days, n=172, p<0.001; Middle, *prde-1* unmated ctrl: 16.1±0.5 days, n=89, *prde-1* mated ctrl: 12.8±0.5 days, n=77, p<0.001; Right, *prde-1* unmated ctrl: 15.3±0.7 days, n=80, *prde-1* mated ctrl: 13.9±0.4 days, n=95, p=0.0333.

**Figure S4.**
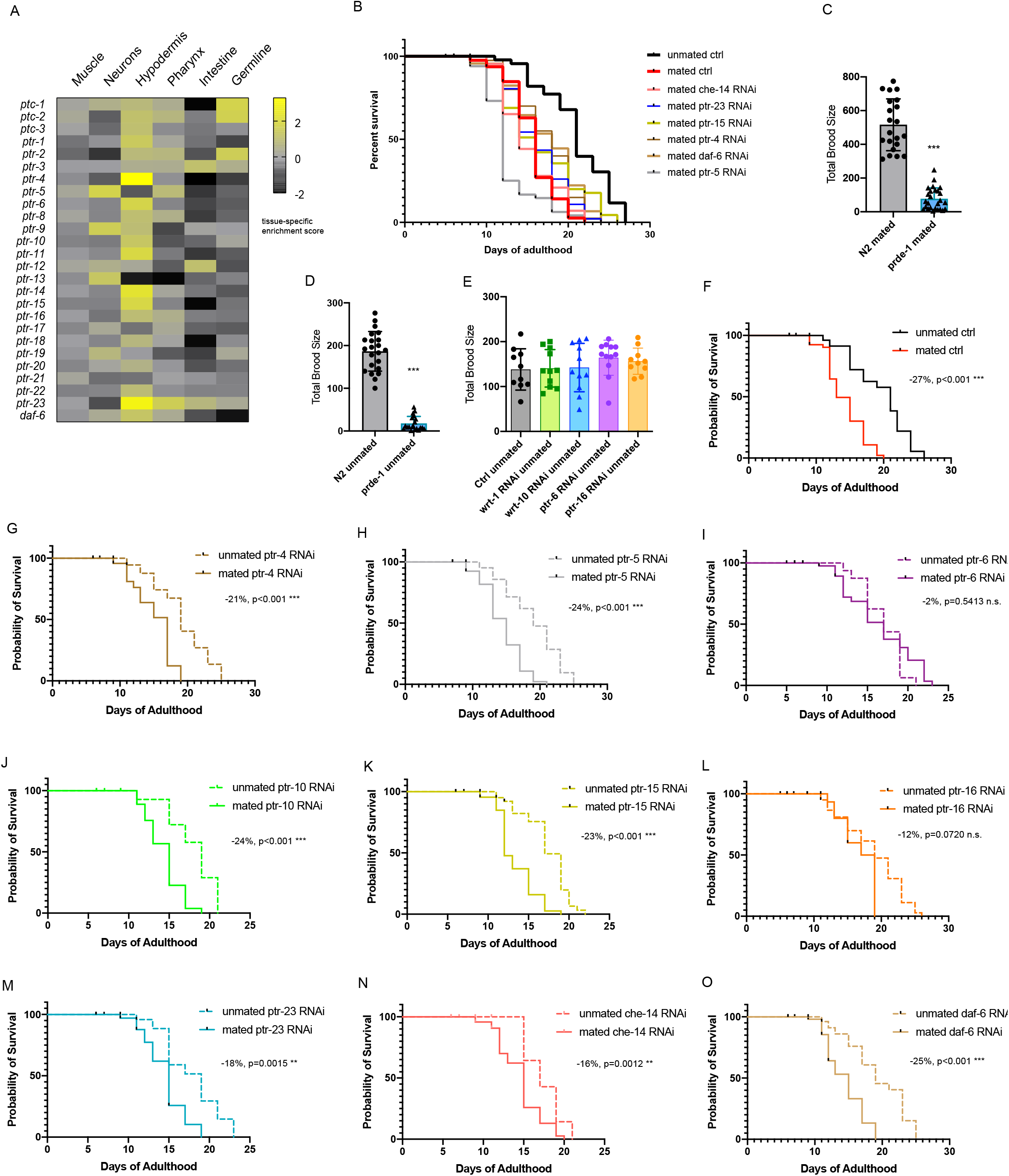
Hypodermal expressing *ptr-6* and *ptr-16* but not other Patched-related receptor encoding genes are required for mating-induced death. A) Tissue-specific expression prediction of Patched receptor homolog genes B) Knock down of Patched receptor-related genes other than *ptr-6* and *ptr-16* fails to rescue the mating-induced death. Mated control: 15.7±0.3 days, n=85; unmated control: 21.1±0.6 days, n=46, p<0.001; mated che-14 RNAi: 15.2±0.5 days, n=46, p=0.7364; mated ptr-23 RNAi: 16.3±0.5 days, n=47, p=0.1611; mated ptr-15 RNAi: 16.7±0.7 days, n=48, p=0.0372; mated ptr-4 RNAi: 17.2±0.6 days, n=49, p=0.0024; mated daf-6 RNAi: 17.3±0.6 days, n=49, p=0.0007; mated ptr-5 RNAi: 12.7±0.5 days, n=53, p<0.001. All lifespans were compared to that of mated worms treated with control RNAi. C) Defective piRNA pathway significantly reduces the total mated brood size. N2 mated: 516±154, n=21; *prde-1* mated: 77±65, n=26, p<0.001, t-test. D) Defective piRNA pathway also significantly reduces the total unmated (self-fertilized) brood size. N2 unmated: 187±46, n=22; *prde-1* unmated: 18±17, n=20, p<0.001, t-test. E) Inhibiting the Hedgehog signaling by RNAi does not affect the unmated (self-fertilized) brood size of the wild-type N2 hermaphrodites. N2 control unmated: 138±46, n=10; wrt-1 RNAi unmated: 141±42, n=11; wrt-10 RNAi unmated: 142±54, n=11; ptr-6 RNAi unmated: 164±39, n=12; ptr-16 RNAi unmated: 156±29, n=10. F-O) Lifespans of mated and unmated *fog-2* hermaphrodites treated with individual Patched receptor homolog genes. Kaplan–Meier analysis with log-rank (Mantel–Cox) test was used to determine statistical significance. Lifespan of mated worms was always compared to unmated worms of the same RNAi treatment. F) unmated control: 19.5±0.9 days, n=53; mated control: 14.2±0.4 days, n=71, p<0.001. G) unmated ptr-4 RNAi: 19.1±0.7 days, n=50; mated ptr-4 RNAi: 15.0±0.5 days, n=67, p<0.001. H) unmated ptr-5 RNAi: 19.0±0.9 days, n=50; mated ptr-5 RNAi: 14.5±0.4 days, n=60, p<0.001. I) unmated ptr-6 RNAi: 16.9±0.6 days, n=40; mated ptr-6 RNAi: 16.5±0.7 days, n=68, p=0.5413. J) unmated ptr-10 RNAi: 18.9±0.6 days, n=49; mated ptr-10 RNAi: 14.3±0.3 days, n=70, p=0.0008. K) unmated ptr-15 RNAi: 17.3±0.5 days, n=70; mated ptr-15 RNAi: 13.4±0.4 days, n=61, p<0.0001. L) unmated ptr-16 RNAi: 18.9±0.7 days, n=55; mated ptr-16 RNAi: 16.7±0.7 days, n=60, p=0.0720. M) unmated ptr-23 RNAi: 17.8±0.9 days, n=50; mated ptr-23 RNAi: 14.6±0.3 days, n=72, p=0.0015. N) unmated che-14 RNAi: 19.4±1.0 days, n=50; mated che-14 RNAi: 14.5±0.4 days, n=64, p<0.001. O) unmated daf-6 RNAi: 17.4±0.6 days, n=50; mated daf-6 RNAi: 14.6±0.4 days, n=69, p=0.0012.

## Tables

**Table 1.** List of significantly differentially expressed genes identified from mated vs unmated *fog-2* germline mRNA_seq

**Table 2.** List of significantly differentially expressed genes identified from mated vs unmated *fog-2* germline smallRNA-seq

**Table 3.** List of germline specific up-regulated genes

**Table 4.** List of germline-specific up-regulated genes that are also up-regulated in mated whole *fog-2* worms with intact germlines

**Table 5.** List of piRNA-targeted germline specific mating-induced genes

